# In silico framework for benchmarking optogenetic hearing restoration

**DOI:** 10.64898/2026.07.13.737798

**Authors:** Lakshay Khurana, Petr Nejedly, Daniel J Jagger, Tobias Moser, Lukasz Jablonski

## Abstract

Cochlear implants (CI) partially restore hearing in profoundly hearing impaired or deaf people by electrically stimulating the auditory nerve. A bottleneck of electrical CIs is the broad spread of electrical current from each electrode that limits the transfer of spectral information, which might be overcome by future spatially confined optogenetic stimulation. Here we established an in silico framework, FraSCO, to model sound encoding in the human cochlea by an optogenetic CI (oCI) for testing the potential of optogenetic hearing restoration. The biophysical modeling framework combined an optical raytracing model implementing a human cochlea implanted with a waveguide-based oCI with a single compartment model of optogenetically modified spiral ganglion neurons (SGNs). The input was an optogenetic sound coding strategy and the quality of the neural representation was evaluated based on comparison of neurograms evoked by optogenetic and electrical stimulation to the spectrogram of the sound applied. The model aimed for technologically feasible properties of the oCIs with 64 stimulation channels. The biophysical modeling framework successfully captured essential physiological features of optogenetic SGN stimulation with a minimal set of ion channel types expressed in the SGN soma. Working with a sample of 1000 SGNs distributed along the tonotopic axis to represent sound encoding, we found that improved spectral selectivity more than compensates for lower temporal fidelity of current implementations of optogenetic stimulation. The established computational framework enables in silico investigation and benchmarking of sound encoding in the cochlea by future oCI and state-of-the-art eCI. The results indicate that optogenetic sound encoding has potential to improve speech understanding in noisy environments for CI users.

## Introduction

Neuroprosthetics, the field at the intersection of neuroscience and engineering, has witnessed remarkable progress in recent decades, empowering individuals with severe neurological conditions to lead more independent and fulfilling lives [1,2]. Cochlear implants (CIs), devices that partially restore hearing for individuals with profound hearing impairment or deafness by directly stimulating the auditory nerve, exemplify this progress. Since their inception in the 1950s, CIs have undergone continuous advancements, with refined sound processing algorithms and intracochlear array designs enabling recipients to perceive speech, music, and environmental sounds with increasing clarity and complexity maximally [3–5].

While the CIs have remarkably improved the lives of its users, these devices also have limitations. The current state-of-the-art electrical CIs (eCIs) stimulate the auditory nerve by delivering electrical pulses inside the electrically conductive medium of the cochlea. This leads to a wide spread of electrical current and it limits the number of perceptually independent electrodes to typically less than 10 [6–10]. This restricts their ability to provide spectral (frequency) information to the brain explaining difficulties of eCI users in understanding speech in noisy environments and appreciating music [3,11–15].

The development of new technologies and sound processing strategies is crucial to address these limitations and further enhance the performance of CIs. One promising approach involves the use of optogenetics [14]. The optogenetic CIs (oCIs), which use light pulses instead of electrical pulses, would offer improved transfer of spectral information as light can be better confined in space when compared to the electrical current [14–18]. This promises improved speech recognition and music appreciation.

Despite good progress and promising preclinical data, oCI development faces several challenges such as the relatively slow kinetics of optogenetic stimulation. The channelrhodopsins (ChRs), used for rendering SGNs light sensitive, do not allow faithful responses to stimulation beyond 200–300 pulses per second (pps) [19–22]. Efforts to optimize ChRs for faster stimulation are underway, but one important question is whether the improved spectral selectivity benefits CI users in terms of speech intelligibility given the currently achievable firing rates? Addressing this question is critical before moving on toward clinical trials on the oCIs. With the framework we present in this paper, we offer an approach that bridges preclinical and future clinical studies by using computational models to predict the intelligibility that could be achieved with oCIs and compare it to the one with eCIs.

Computational models have been used in the development of CIs since the 1970s as a powerful tool for understanding the underlying mechanisms of hearing restoration and to accelerate the development of more effective implant designs and sound coding strategies (for reviews of computational models in CI research see ref. 23–25). These models aim to recapitulate the function of the auditory system at varying depth, allowing researchers to investigate various aspects of CI operation, such as electrode-neural interface, neural encoding on the predicted outcome performance and individual outcome variation.

By combining the stage-specific models with sound coding strategies, computational frameworks can be formed to investigate broad research topics, as has been done to predict speech intelligibility [26–29]. In this paper, we present a modular framework to predict information transmission from sound to spikes for optical and electrical stimulation called FraSCO (framework for sound coding optimization). The framework includes simulation of CI sound coding strategies, prediction of spatial spread of current and light, a single-compartment SGN model including voltage-gated channels and a ChR for optogenetic activation, and a similarity index to compare the information in spikes to the sound spectrogram.

## Methods

### Framework structure

A modular framework structure with the stages of (1) sound processing, (2) optical or electrical stimulation, (3) spiral ganglion neurons, and (4) analysis of neural code. (Figure 1; each stage is described in the following sections) was implemented in MATLAB R2021a (The MathWorks Inc.), unless stated otherwise.

**Figure 1.**
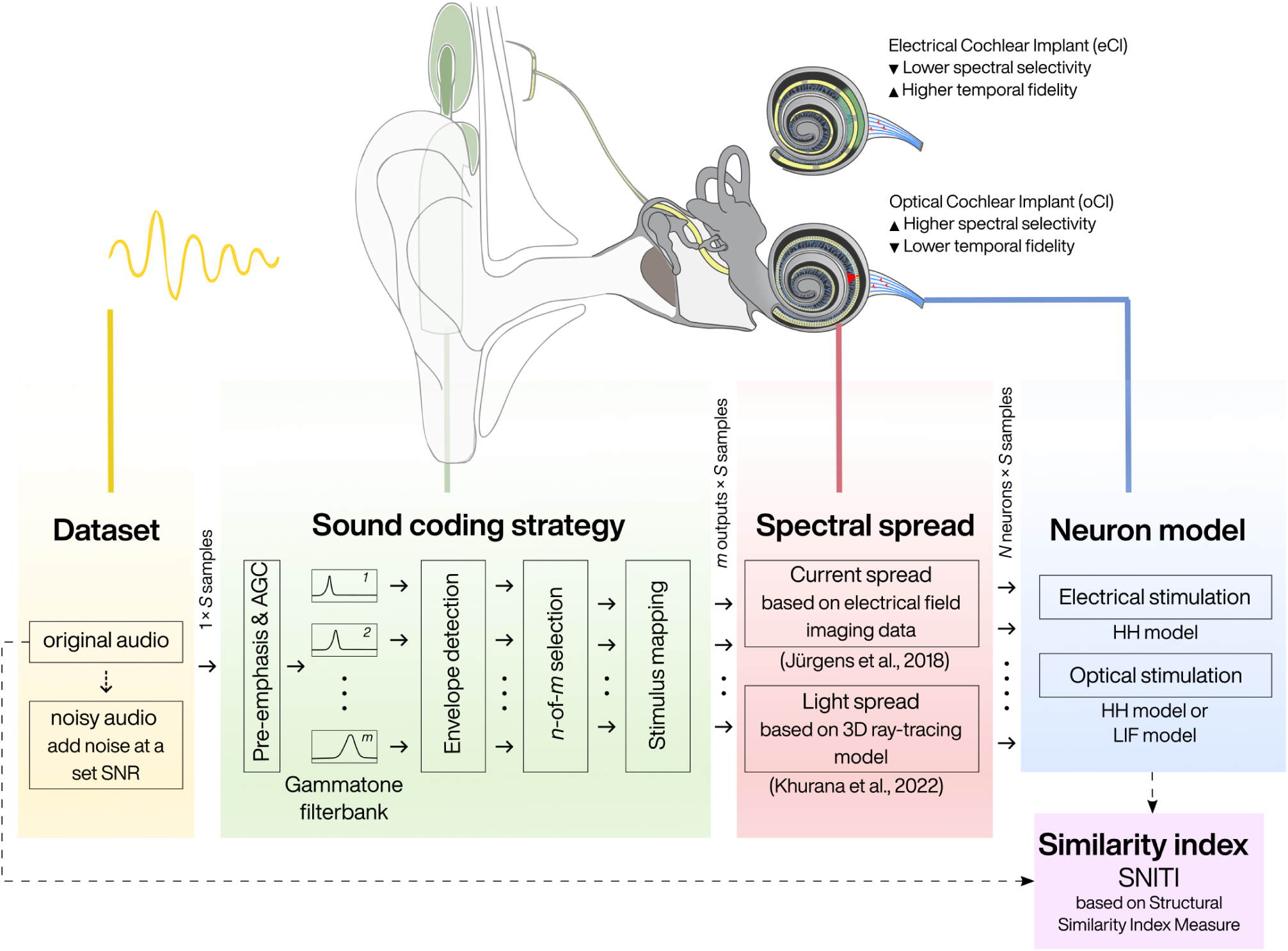
Overview on the complete framework. An audio file is read from a dataset and resampled at 100 kHz. Either the clean signal or a noisy signal (at a set SNR) serves as input to the sound coding strategy block. Here, the signal is sampled in frames and processed with n-of-m strategy, with *m* total channels out of which *n* active channels with highest energy in the frame are selected for stimulation. The electrodogram/emittogram output of coding strategy block determines how the current/light spreads in the cochlea along the tonotopic axis using the Gaussian spread values from Jürgens et al. (2018) [37] and Khurana et al. (2022) [15]. The output is a matrix of current/light values at the spiral ganglion neurons. The SGN model is an extended Hodgkin-Huxley model with Markov kinetics formulation. The ChR model is integrated in the SGN model for optical stimulation. Additionally, a simpler and faster, Leaky Integrate-and-Fire model is implemented for the optical stimulation with higher spike probabilities. The output of the 1000 SGNs composes a spike raster plot evoked by optical or electrical stimulation. Finally, a similarity index is computed by comparing the spike intensity and spectrogram from the clean audio. The framework is modular and any block can be replaced with a different model, for example, a new sound coding strategy could be implemented.

### Sound processing and stimulation strategy: the n-of-m strategy

Any implementation of a sound coding strategy, either a clinical, experimental, or conceptual one, can be used in this stage block. We implemented a generic eCI strategy, *n*-of-*m* (the generic form of the Advanced Combined Encoder, ACE, strategy [30]), which forms the basis of most clinical strategies currently in use. All parameters, necessary for testing a variety of inputs, such as pulse rate, pulse width, total number of channels (*m*), maximum number of channels stimulated per frame (*n*), and maximum stimulus amplitude can be varied in the implementation.

The strategy (1) samples the audio in frames, (2) applies a pre-emphasis filter, (3) filters the signal through *m* number of infinite impulse response gammatone filters, (4) calculates the energy in all filter outputs, (5) selects the maximum *n* filters with highest energies, (6) maps the energy using loudness growth function, threshold and comfort levels, and (7) generates the electrodogram, in case of eCIs, and emittogram, in case of oCIs.

### Cochlear stimulation by current and light

The output values (*z*) of the sound processing which ranged from 0 to 255 was mapped to amplitudes (*A*) using the equation

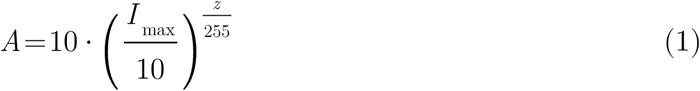

where the maximum intensity (*I*_max_) was set for current to 1750 µA and for light to 200 mW for the simulations presented.

The spread of current and light from the source to the neurons was estimated using already published data [15]. An optical ray-tracing model, implemented in TracePro Standard 7.8.1 (Lambda Research Corporation), predicted the spread of light using three different light sources, namely, LEDs, Laser-coupled waveguides with numerical aperture (NA) 0.5 and 0.17, in a human cochlea model. We had previously compared the spatial spread of light with the current spread [15] and provided a detailed description of the model and measurement of the spatial spread of stimulation. Here, we used the median values of the spread of light from oCI waveguide outcoupling structures (NA 0.5) and eCI electrodes, and calculated spread at 1000 neurons across the tonotopic axis. We note that the framework can readily accommodate other sets of spatial spread of stimulation for evaluation of the effect of different light emitter properties or various electrode configurations.

### Biophysical single compartment model of SGNs

We developed an extended Hodgkin-Huxley (HH) model using Markov kinetic formalism in Python 3.8 programming language. The SGN model is described as

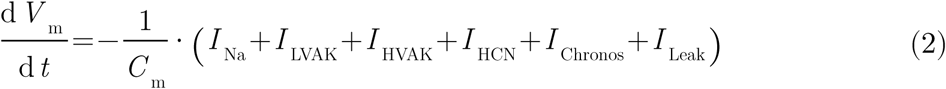

where the *I*_Na_, *I*_LVAK_, *I*_HVAK_, and *I*_HCN_ are the individual ion currents of voltage-gated sodium (Na), low-voltage activated potassium (LVAK), high-voltage activated potassium (HVAK), and hyperpolarization-activated cyclic nucleotide-gated (HCN) channels, respectively. Additionally, *I*_Chronos_ is the current from the Chronos model for optogenetic stimulation [31]. These ion currents are computed for an ion channel *x* as:

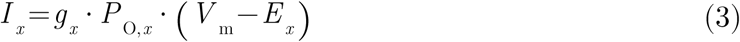

where *g_x_* represents the maximal conductance of the ion channel, *E_x_* is the channel reversal potential, and *P*_O,*x*_ is the probability of being in open state. This probability is computed using Markov models.

The sodium channel consisted of six states—four closed (C), one open (O), and one inactivated (I)—as described by Tveito and Lines (2016) [32]. The equations for transition rate parameters were adopted from the same literature source. Similarly, LVAK, HVAK, and HCN were modelled as CCCCOI (without a loop), CCCCO, and CO Markov models. The models and equations are provided as Supplementary Information (Supplementary Figure 2). In order to fit the parameters of the channel models to the currents of SGNs, we used patch-clamp recordings from SGNs isolated from the mouse cochlea around the onset of hearing described in Smith et al. (2015) [33] and Browne et al. (2017) [34]. Fitting employed an optimization routine, consisting of differential evolution genetic algorithm and leave-one-out cross-validation. As the recordings were conducted at room temperature (22– 24°C) and we run the model for body temperature (37°C), the transition rates were multiplied with respective *Q* _10_ factors (Supplementary Table 1). The open-state probability *P*_O,*x*_ in Equation 3 was obtained as the fraction of channels in the open (O) state of the Markov models. More information about the model parameterization and the *Q* _10_ factors is provided in the Supplementary Information.

The biophysical HH model described here was designed to accommodate both electrical and optical stimulation. However, to achieve higher spike probabilities specific to optogenetic stimulation, we developed an additional Leaky Integrate-and-Fire (LIF) model tailored for this purpose [35].

### Analysis of neural code: Sound-to-Neuron Information Transmission Index

In order to represent the neural response to oCI or eCI stimulation, the spike raster plot was converted to a spike intensity plot by assigning a value to each element of the matrix equal to the average spikes in a small window centered at that element. This was mathematically described as

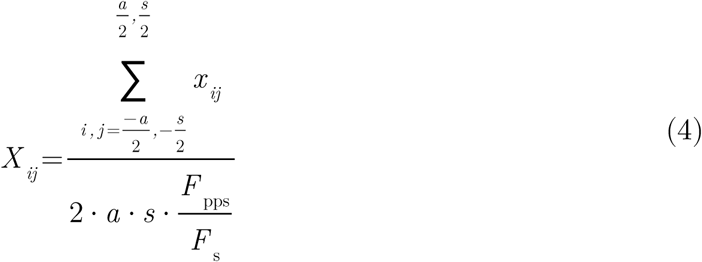

where *x* is the spike raster plot and *X* is the spike intensity matrix with values obtained by averaging the window of *a* neurons and *s* samples, *F*_pps_ is the stimulation rate, and *F*_s_ is the sampling frequency.

The spectrogram of the input audio (*S*) was then compared to this matrix (*X*) using the inbuilt function in MATLAB for the Structural Similarity Index measure (SSIM). It computes a score on various windows using

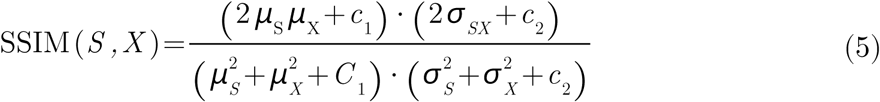

The SSIM score is a good measure to compare different parameter sets of processing the same input audio. However, this raw similarity score reflects both the signal and silence similarities. Silence within an audio file can artificially inflate similarity scores, leading to biased results that do not accurately reflect the true similarity based on the signal of the audio. To isolate the contribution of silence, a reference score was calculated for each file using a raster plot with no evoked spikes (*X* _0_). The final score was fit on a sigmoid curve centered around the reference score, which is the score when there is no stimulation, effectively removing the bias introduced by varying amounts of silence, yielding a normalized score that more accurately reflects the true content similarity. We termed this measure “Sound-to-Neuron Information Transmission Index” (SNITI), and it is computed as

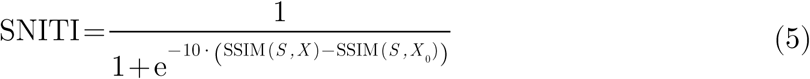

SNITI score of 0.5 represents no stimulation. Higher values indicate higher similarity between input audio and sound coding output. Lower values indicate more noise than the input audio.

Dataset

A speech corpus [36] containing audio files of single words (such as “character”, “glass”, “strategy”, “plant”, and “thin”), was used for the analysis described in the section Eexmplary applications of the framework. The corpus consists of 50 words from a male speaker and 50 words uttered by a female speaker, out of which 20 files were selected at random from each speaker. Silent beginning and end longer than 10 ms were clipped to save processing time. The corpus contains high-quality (256 kbps bitrate) recordings at a high sampling rate of 48 kHz with very low background noise. We resampled the audio files at 100 kHz.

Statistical analysis

To compare the performance of eCI and oCI across various stimulation rates, a two-way ANOVA was performed. The two factors considered were the stimulation rate and stimulation type (electrical or optical). The interaction between these two factors was also evaluated. The data were normally distributed, and the assumptions of ANOVA were checked and met.

Following the two-way ANOVA, a post hoc analysis was performed using the in-built ‘multcompare’ function of MATLAB to examine pairwise comparisons across both dimensions, comparing eCI and oCI performance across all stimulation rates.

## Results

The framework of predicting sound to neuron transmission of information

To predict the transmission of auditory information from sound to neural firing we developed a modular framework for modeling optical as well as electrical SGN stimulation (Figure 1). The input audio is transformed by the sound processing stage to generate a sequence of current or light pulses which are spread-out spatially along the tonotopic axis (i.e. in the frequency domain). SGN somata located in the Rosenthal’s canal of the cochlear modiolus respond to the stimulation and the ensuing spike patterns are used to compare the activity with the spectrogram of the input audio. This comparison is done using a new measure SNITI—Sound-to-Neuron Information Transmission Index (see Methods)—based on Structural Similarity Index Measure (SSIM).

### Sound processing

All sub-blocks are identical for electrical and optical stimulation except for the pulse shape. Biphasic pulses of 60 µs per phase are generated for electrical stimulation, whereas the optical stimulation uses monophasic pulses of 1 ms. A comparison of the output of sound processing for electrical and optical stimulation is provided in Supplementary Information (Supplementary Figure 3).

### Response to electrical and optical stimulation

The HH SGN model (Figure 2 A) was assembled from 4 different channel types: voltagegated sodium channels (NaV), low-voltage and high-voltage activated potassium channels (LVAK and HVAK, respectively), as well as hyperpolarization and cyclic nucleotide-activated channels (HCN), as described in the respective Methods subsection. Figure 2 B exemplifies the model by showing the fits of the NaV channel model to NaV currents recorded from mouse SGNs [34] and the plots for other channels are provided in Supplementary Figure 2. Action potential firing of the SGN was fitted to the extended experimental data from Rutherford et al. (2012) [38]to optimize the remaining parameters of conductance and capacitance. The model’s response was well-correlated with the recorded traces, as demonstrated in the examples in Figure 2 C for increasing ramp stimuli with different maximum amplitudes and different durations. Finally, an existing 8-state Markov model of Chronos [31] was integrated to the SGN model to drive spiking by optogenetic stimulation with the fastest native ChR reported thus far [21,39]. To validate the optical stimulation of the model, we compared the spike probabilities at various stimulation rates of the modeled neuron with the average spike probabilities recorded by Keppeler et al. [21] (Figure 2 D). The comparison revealed a strong correlation, with both the HH model and the recorded data showing a gradual decrease in spike probability as the pulse rate increased from 20 pps to 900 pps. Additionally, a Leaky Integrate-and-Fire (LIF) neuron model was implemented with higher spike probabilities which captured the firing of the fastest SGN response recorded with Chronos [21]. This approach allowed us to do the simulations for the currently achievable average light-driven firing behavior of SGNs, as well as for faster SGN firing that are realistic and attainable with future advancements.

**Figure 2.**
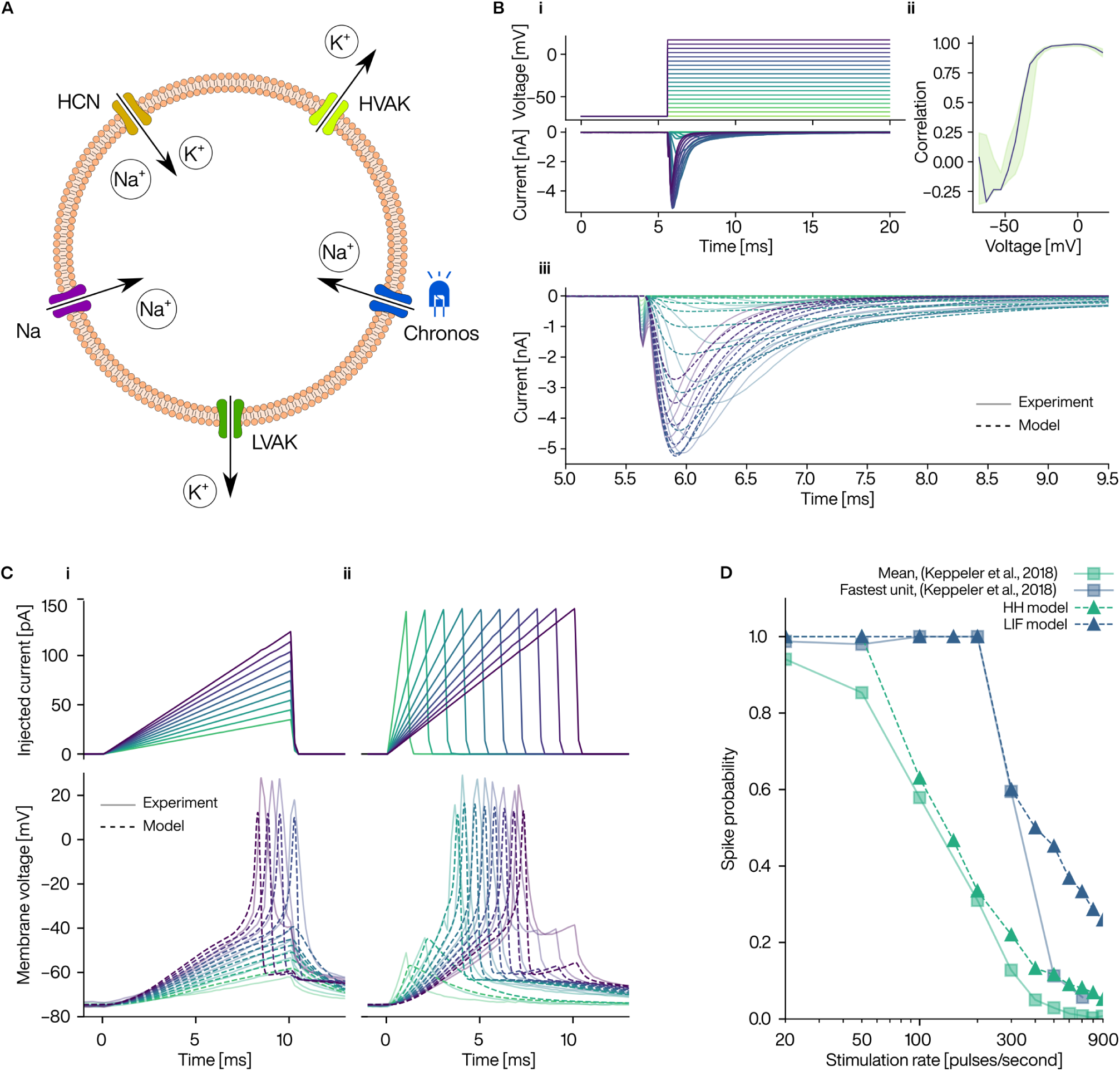
Development and behavior of the SGN model for electrical and optical stimulation. (A) Illustration of the single-compartment Hodgkin-Huxley (HH) model of the SGN. The model consists of voltage-gated sodium (Na_V_), low-voltage activated potassium (LVAK), high-voltage activated potassium (HVAK), and hyperpolarization-activated and cyclic nucleotide–gated (HCN) channels. In addition, a channelrhodopsin, Chronos, was added for optical activation. (B) Example fitting of Na_V_ channel parameters using Markov kinetics formalism. The Markov model and equations are described in Supplementary Information (Supplementary Figure 2). (i) Voltage clamp protocol (top) and corresponding recorded data (bottom) calculated as mean from ten cells. (ii) Correlation of recorded data from the experiment and response of the model. Black line corresponds to the final model. The green area shows maximum and minimum correlations from all leave-one-out cross-validations. (iii) Response of the model (dashed lines) for the same voltage clamp protocol as shown in (i), overlaid on the recorded data (faded lines). (C) Example fitting of the complete SGN model for electrical stimulation. Current clamp protocol (top) of increasing ramp stimuli with (i) changing maximum amplitudes and (ii) changing durations, and responses (bottom) from the recorded SGN (faded lines) and the model (dashed lines). The timepoint of zero indicates the onset of the ramp. (D) Spike probability for optical stimulation of the SGN for increasing pulse rate. The model data are compared to the published data [21]. The spike probabilities of HH model show a correlation to the mean from the recordings of 40 putative SGNs (obtained from 6 mice). The spike rate of the fastest unit recorded was 100% at 200 pulses/second and decreased to ∼60% at 300 pulses/second. The Leaky Integrate-and-Fire (LIF) model follows this pattern of spike probability. The stimuli for the model simulation had 1 ms long rectangular pulses at an intensity of 100 mW.

### Comparison of input and output

In this final step, the spike raster plot generated by the neuron model is compared to the spectrogram of the input audio. Several studies have previously established that SSIM, which was originally designed to compare similarity of two images, can be used to compare two neurograms and to predict speech intelligibility [40–43]. The SSIM or SSIM-derived measures, such as, Neurogram Similarity Index measure (NSIM), compare one ‘clean’ or ‘original’ neurogram to a ‘degraded’ or ‘processed’ neurogram. In our case, we would like to compare two processed raster plots, which contains only binary values, with the spectrogram of input audio. To do so, we used the SNITI measure described in the Methods section.

When there is no stimulation, a score 0.5 would be achieved. Higher values indicate higher similarity to the input audio and lower score mean more noise than the input audio. A value of 0.5 can also be obtained with stimulation when the amount of ‘similar’ information is balanced by the amount of noise. The SNITI scores may not linearly correlate with subjective intelligibility scores, however, we are using the measure for relative comparison only.

Exemplary applications of the framework

We employed the framework to investigate the effect of the number of stimulation sites, the stimulation rate, and noise on SNITI scores for electrical and optical stimulation. For the description of dataset used for these simulations, please see the Methods section.

### Effect of the number of stimulation sites

The total number, *m*, was limited to a maximum of 20 for eCI because of the interleaved stimulation at high pulse rate with pulse width of 60 µs, whereas, parallel stimulation of oCI allowed high number of emitters that we scaled up to 64 emitters. The total number of channels (*m*, electrodes in case of eCI and emitters in case of oCI) was gradually increased and the number of active electrodes/emitters (*n*) was varied from 4 to 16 for the eCI and up to 64 for the oCI. The pulse rate was set to 900 pps for eCI and 150 pps for oCI (HH model). The SNITI score increased for eCI with increased *m*, but for a fixed *m*, the score did not show improvement above 10 active channels, saturating around 0.75 score (Figure 3 A). This correlates with observations in eCI users with the most eCI configuration (mid-scala electrode, monopolar stimulation) who do not show improvement in speech understanding above 8 or 10 channels [6–10]. The scores for oCI for the same *n* and *m* pairs were lower than for eCI, which likely reflects the lower stimulation rate of 150 pps. When the total number of channels were increased for oCI, the SNITI scores improved and reached a maximum of 0.71 for 48-of-64 configuration (Figure 3 B).

**Figure 3.**
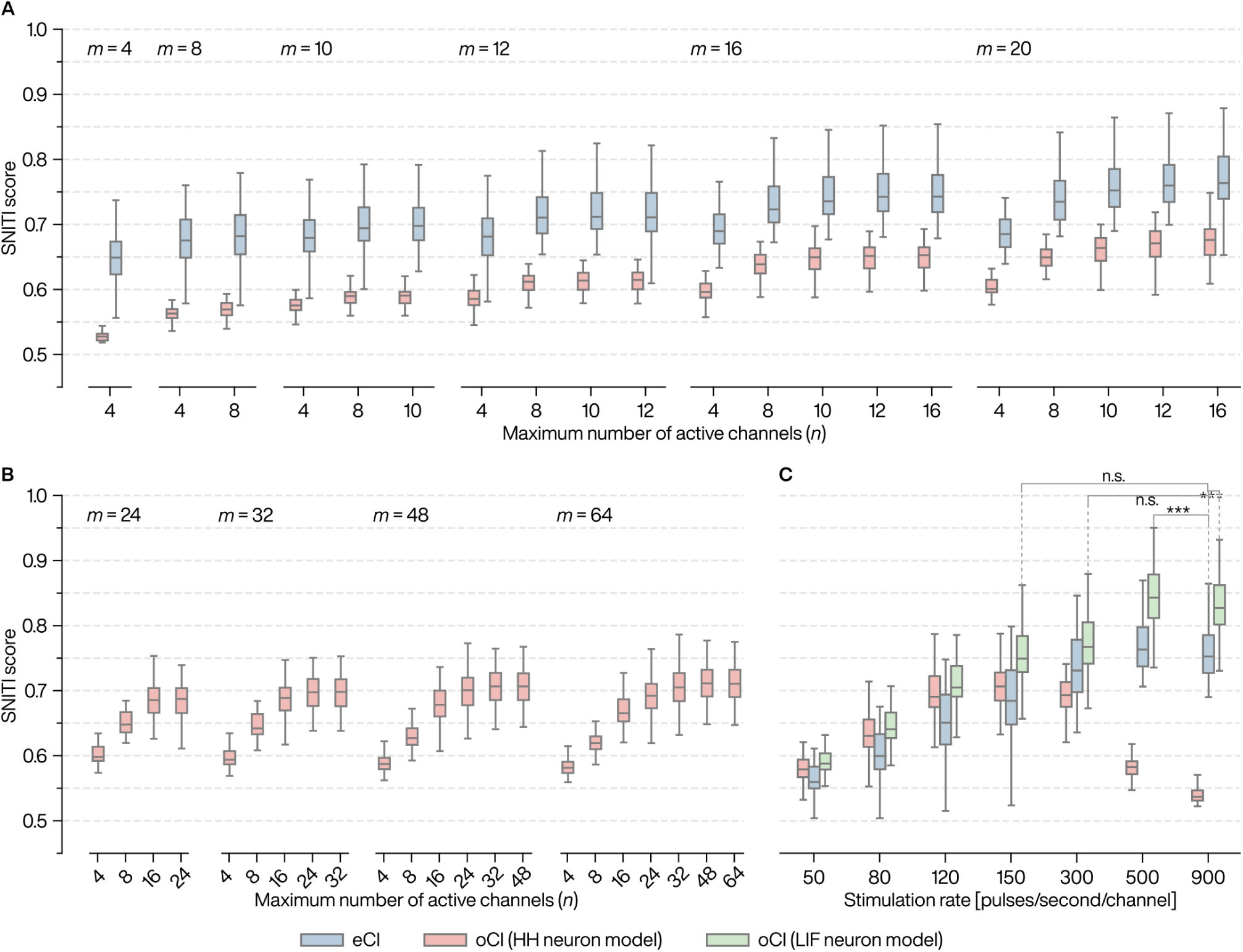
Effect of the total number of stimulation sites (m), maximum number of stimulation sites per frame (n), and stimulation rate on similarity index. (A) Sound-to-Neuron Information Transmission Index (SNITI) scores as obtained from the framework for a dataset of 40 audio files of single words, with various combinations of number of stimulation sites (*m*) and number of maximum active channels per frame (*n*). The same sound processing parameters were used for the simulation of eCI (in blue) and oCI (in red), except that the stimulation rate and pulse duration were 900 pps and 60 us for eCI and 90 pps and 1 ms for oCI. (B) SNITI scores for extended combination sets of *n* and *m* for oCIs as the parallel/simultaneous stimulation of multiple light emitters offers greater number of *m*. (C) SNITI scores for increasing stimulation rate of eCI (in blue), oCI with HH neuron model (in red), and oCI with LIF neuron model (in green). The *n*-of-*m* was fixed to 10-of-20 for eCI and 32-of-64 for oCI. Statistical significance (*** for p<0.001) or non-significance (n.s.) is indicated only for selected post hoc comparisons, namely, the eCI at 900 pps with certain oCI configurations of the LIF model. The absence of a significance marker does not imply a lack of significance or relevance. Also, the scores for oCI with LIF model were significantly higher than eCI for the same stimulation rates above 50 pps. Each box plot shows the median, upper quartile and lower quartile. The whiskers extend to the minimum and maximum values excluding the outliers (not shown).

### A potentially lower stimulation rate in oCI can be traded-off with higher spectral resolution

The effect of stimulation rate was evaluated at pulse rates of 50, 80, 120, 150, 300, 500 and 900 pps (Figure 3 C). The *n*-of-*m* were fixed to 10-of-20 for eCI and 32-of-64 for oCI.

We found that the median score of oCI with HH model was highest at 150 pps and the scores did not show a significant improvement over eCI scores. With the LIF model, we expanded the predictive capacity of our framework with the SGNs capable of more faithful stimulation at rates beyond 150 pps, as observed in some in-vivo recordings [21]. At low spike rates, the LIF model and the HH model exhibited the same behavior in terms of SNITI scores. However, due to greater spike probabilities at higher pulse rates, oCI stimulation results with the LIF model outperformed not only the scores with HH model as well as the eCI scores. The two-way ANOVA between eCI and oCI (LIF model) revealed significant effects for both the stimulation rate (p<0.001) and stimulation type (p<0.001). Additionally, the interaction between stimulation rate and implant type was significant (p<0.01), indicating that the effect of stimulation rate on performance varied depending on the type of cochlear implant. Post hoc comparisons revealed that the oCI consistently outperformed eCI across all stimulation rates (p<0.01 for 100 pps, p<0.001 for 120, 150, 500 and 900 pps, p<0.05 for 300 pps) except for 50 pps. Notably, the performance of eCI at the highest stimulation rate of 900 pps did not differ significantly from that of oCI at 150 pps or 300 pps. However, eCI at 900 pps performed significantly worse than oCI at 500 pps (p<0.001) and at 900 pps (p<0.001). Additionally, in agreement with the literature on psychoacoustic tests with eCI users, where the score typically plateaus beyond 300 pps except for a few subjects [9,44–46], the simulated eCI performance at 900 pps had no significant differences to that of 300 or 500 pps.

While clinical eCIs operate at higher pulse rates but achieve spike probabilities of only about 10% to 20% [47], optically activated SGNs can attain spike probabilities as high as 60% at a stimulation rate of 300 pps [21], resulting in firing rates comparable to the electrical stimulation at 900 pps. Moreover, the improved spectral selectivity of oCIs likely contributes to the enhanced performance observed in simulations with the LIF model.

### Noise impacts oCIs less drastically than eCIs

It is important to note that the dataset used for these simulations is high-quality and almost noiseless. This can be equated with psychoacoustic testing of CI users in a quiet lab environment, where they are known to achieve high understanding of speech. Real-world situations are rarely without any noise, and eCI users struggle with understanding speech in noisy environments. Therefore, we evaluated the effect of added noise and compared it for eCI and oCI (with the HH model; Figure 4). White noise was added to each audio file in the dataset using the in-built function, ‘awgn’, of MATLAB. The signal-to-noise ratio (SNR) was gradually decreased from +20 dB to −20 dB. Median SNITI scores were calculated and a linear fit was done to find the slope. The scores decreased at a rate of 0.017 dB^-1^ for eCI, whereas for oCI the decrease at 0.009 dB^-1^ was almost half of the slope for eCI. In light of these results, potential oCI users are predicted to be less impacted with noisy backgrounds, already with the state-of-the-art opsins, and would will likely experience further improvement with optimized ChR and more sophisticated sound coding strategies.

**Figure 4.**
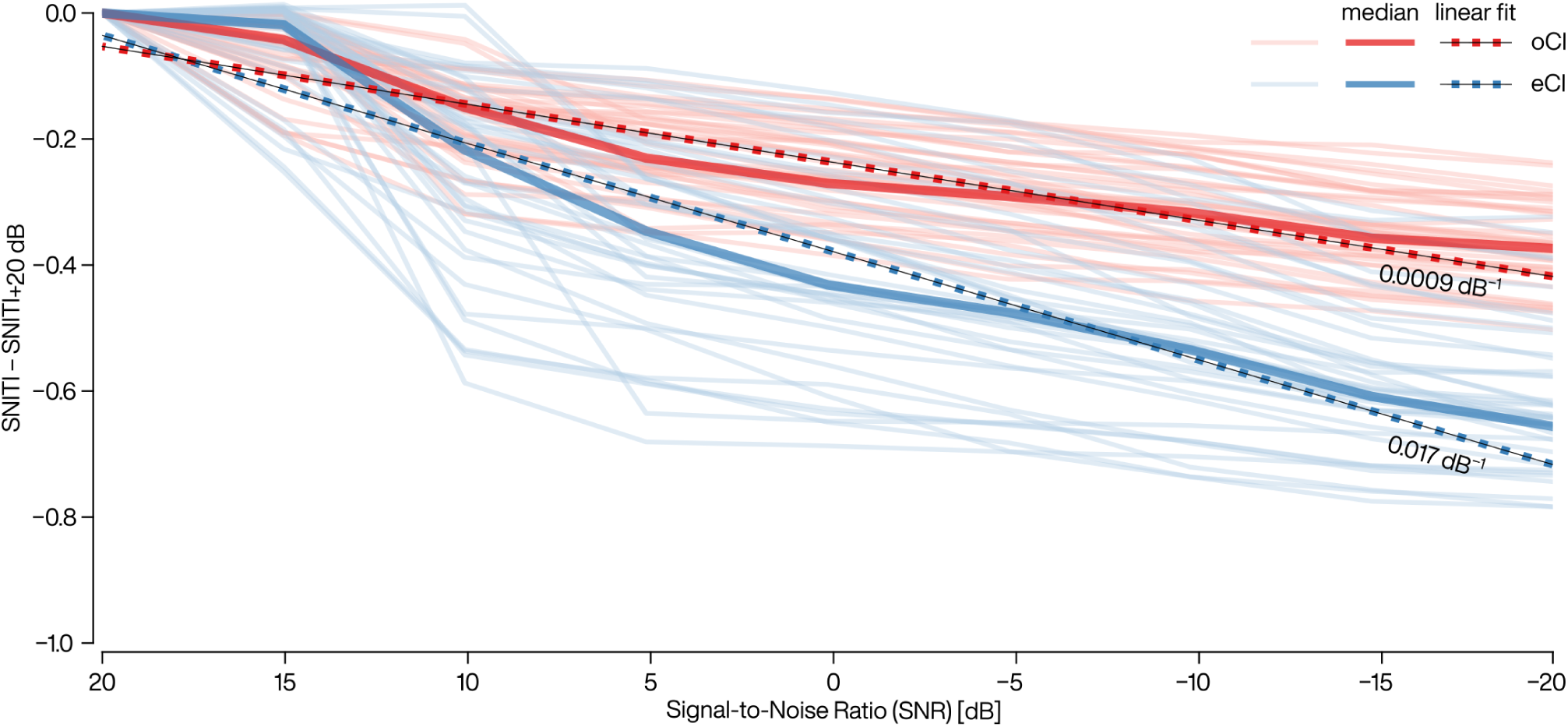
Effect of changing Signal-to-Noise Ratio (SNR) on similarity index. White noise was systematically added to each audio file at SNR from +20 to –20 dB, in steps of 5 dB. Each noisy file was analyzed with the framework to compute the similarity index, SNITI. Data points for each file are plotted as a connected line for eCI (in light blue) and for oCI with the HH model (in light red). The values were adjusted to start the line at zero on y-axis for easy visual comparison. The best linear fit on the median had a slope of 0.017 and 0.009 dB^-1^ for eCI and oCI, respectively.

## Discussion

The major limitation of eCIs is the low number of perceptually different channels, due to the wide spread of current in the cochlea [6–10]. Studies with electrophysiological recordings in the inferior colliculus of mouse [14,16] and gerbil [17,18] and computational models [15] have previously shown that the spectral spread of excitation is much lower for optical stimulation than electrical stimulation. But how this improvement would translate to speech understanding, remained unanswered. Here, we have presented a computational framework for prediction and comparison of speech information at the level of auditory nerve fibers after stimulation with eCI and oCI. The framework is a chain of modular blocks of CI sound processing, current and light spread, SGN response, and a similarity index. The sound processing block was developed based on generic *n*-of-*m* coding strategy. The spatial spread of excitation was calculated using existing models and data. The SGN response was predicted using a biophysical model of optogenetically modified SGNs, constructed by fitting ion-channel models to electrophysiological recordings. The information of the neural firing across the tonotopic range was compared to the sound spectrogram using a similarity index, SNITI, based on an existing similarity measure, SSIM. Using the framework, we compared the SNITI scores for eCIs and oCIs for a variety of number of total channels and number of maximum active channels in a frame of stimulation, the stimulation rate, and the addition of noise.

Several modelling efforts have been presented in the past to answer a variety of questions related to the eCIs [23,26–29,48,49], however, up to our knowledge, our framework is the first one to simulate electrical as well as optogenetic stimulation in the same model framework. The major benefit of the framework would be to assist the development of oCIs by narrowing down the parameters of the light sources and by guiding the development of opsins. Moreover, oCI-specific sound coding strategies could be investigated using the presented framework.

### Framework application

As an application of the framework, we simulated the eCI and varied the number of channels to reproduce the limitation of perceptually different channels. We found no improvement in the similarity measure for maximum number of channels above 10 for eCI, in agreement with the psychophysical studies with eCI users [6–10]. The scores for oCI, however, continued to gradually increase up to 48 channels, demonstrating the advantage of higher spectral resolution. This advantage is clearer when comparing the electrical and optical stimulation at different stimulation rates. The oCI, modeled with attainable fast stimulation, with higher spectral resolution (32 active channels) performed significantly better than eCI at same stimulation rates. Also, oCI (LIF model) at stimulation rates of 150 and 300 pps achieved comparable performance to eCI at 900 pps, indicating the trade-off between the spectral and temporal resolution of the stimulation. One reason for this tradeoff, other than the increased spectral selectivity of the oCI, is the comparable firing rates of oCI at 300 pps (with ∼60% spike probability for the LIF model) and of eCI at 900 pps (with ∼20% spike probability [47]). Although the promising results of the simulations are based on the neuron model characterized by fast dynamics (LIF neuron model), it is reasonable to predict that these outcomes could be achieved and further optimized with the development of improved opsins that offer even faster temporal fidelity and better transduction. Recordings of optogenetically driven firing from single putative SGNs suggest that Chronos, the fastest naturally occurring ChR, as well as f-Chrimson and vf-Chrimson do not enable reliable spiking at 300 Hz (average spike probability of 10–20% [19,21,22]). We reason that non-optimal membrane targeting, low single channel conductance (probably around 40 fS for all three ChRs), closing kinetics (from 0.8 ms for Chronos to 3 ms for f-Chrimson) and recovery of light-driven SGN firing during the interstimulus interval [22] all contribute to limit the temporal fidelity of optogenetic sound encoding. Additionally, the integration of a dedicated sound coding strategy tailored to the unique properties of optogenetic stimulation could enhance performance, potentially making optical stimulation an even more viable alternative to conventional eCIs.

The high light intensity threshold of the opsins in another crucial factor hindering the development of oCIs. In our model, the light sources had a maximum power of 200 mW, which is currently a challenge for laser, waveguide, and battery technologies to build such light sources with high power efficiency and small form factor. In recent studies, light probes have been fabricated using flexible waveguides with output power of up to 13 mW and 33% optical efficiency with 20 µm outcoupling structure [50], and using vertical-cavity surface emitting lasers with up to 50 mW power for 450 µm long semiconductor chip [51]. These light intensities are far below the intensity used in our simulations. While the future iterations of the SGN model with more realistic channel expression and multiple compartments may show a reduction in light intensity requirements, nevertheless, developing ChRs conveying higher light sensitivity to SGN firing and improving optical stimulation technologies remain important objectives. This could, for example, partially be tackled with a stimulation strategy based on ultrashort pulselets which are integrated by the SGNs [22]. A discussion on the energy budget of oCIs has been presented in ref. 52.

Background noises interfere with speech and make it challenging for CI users to filter the useful information. While modern CIs have noise reduction algorithms which may show an improvement in speech understanding in noise in a controlled, lab environment, they do not benefit from this in daily-life situations [53–55] and some aggressive noise suppression method may even have detrimental effect on listening when the target sound is identified as noise [55]. The speech understanding, naturally, decreases with decrease of SNR for CI users; however, our framework predicted this degradation to be greater for eCI than oCI. The reason behind this effect could be the combination of two factors—greater number of stimulation sites and narrower spread of stimulation than eCI—which allows recruitment of more stimulation sites where useful information is transmitted and less spread of noise compared [56]. Additionally, the richness and complexities of music cannot be fully captured and transmitted to the SGN with eCIs; therefore, another potential benefit of high spectral resolution of oCIs would be the music appreciation, which was not evaluated in this study.

### Limitations and outlook

The framework presented here aimed to model an average behavior of eCI and oCI stimulation. The parameters of the models approximate the vast complexity of the system. Although adding more complexities require making more assumptions, certain sections of the framework can nevertheless be “upgraded”.

We implemented a generic sound coding strategy (n-of-m) which could be replaced with a specific clinical CI strategy, such as, HiRes120 which uses current steering to create virtual channels between the physical electrode contacts (however, this would only change the focus of stimulation, not the spatial spread). Due to its modular design, our framework serves as a tool for the further studies on the development and optimization of such coding strategies.

The computational models rely on various assumptions and approximations to simplify the simulations of the vastly complex nature of the biological system and the interactions between light/current and tissue. For example, (a) the neuron locations and density were not obtained from imaging data, rather the neurons were placed equidistantly along the centerline of the RC, however, their density is non-uniform across different regions of the cochlea with highest density in the middle region [57], (b) an “average” model of Type-I SGNs was designed whereas the Type-I SGNs have different subpopulations of high, medium or low spontaneous discharge rate [58,59], (c) the Type-II SGNs—forming 5–10% of the SGN population [60,61]—were not included in the model, (d) the current interactions between the electrode contacts were not considered, only the spread was computed, most likely resulting in higher similarity scores, and (e) a simplified point-like SGN model was developed, potentially limiting the ability to capture its full complexity.

Moreover, the whole framework relies on the parameter estimates from experimental studies, which may be subject to uncertainty and variability. For instance, (a) estimates of light scattering and absorption coefficients may vary across different measurement setups, (b) estimates of ion channels’ kinetics and opsin expression levels may vary across different experimental conditions and cells, and (c) neural response thresholds may vary across different stimulation protocols and animal models. All these factors lead to uncertainty in the predictions of the computational models.

Although our neuron model implementation takes advantage of vectorization and multithreading processing, the HH model still has a major impact on computation time (Supplementary Figure 3). A more efficient method could be implemented by either using a phenomenological back-end to the biophysical model [28] or by training a convolutional neural network for a variety of stimuli.

Furthermore, we proposed the similarity index, SNITI, for relative qualitative comparison of different parameter sets in the framework. To transfer the SNITI scores to intelligibility scores, clinical evaluation could be done by using the same dataset and parameters with eCI users. For the evaluation of speech-in-noise, we used white noise, but in daily life, CI users would come across different noises, such as factory floor, classroom, cafeteria, steady state, and multi-talker babble noise. While, these could be evaluated in the framework, the SNITI score might not be a good measure for all noise types. Especially for the noises which overlap in the frequency domain with the target sound, the SNITI measure—which looks for correlation with sound spectrograph rather than intelligible information—could categorize some noise as target because the SGN spikes that resulted from noise might match the sound spectrograph.

## Conclusions

A computational framework was constructed by assembling the models for processes from sound to spikes to predict the similarity in the sound and SGN activity for stimulation with eCI and oCI. The proposed framework could be used to investigate questions and evaluate parameters of sound processing strategies, ChR development and medical device development, as well as, for prediction of relative speech understanding for such parameters. The framework is not limited by the practical constraints of technological limitations and clinical studies. The applications presented in this study predicted that even when assuming a lower temporal fidelity of sound coding in oCIs this is balanced by the higher spectral resolution, even without a dedicated sound coding strategy which can maximize this benefit. In addition, the simulations predicted lower impact of noise in oCIs than in eCIs. In future work, the framework can be expanded for more realistic models and stimuli.

## Acknowledgements

The authors would like to thank Dr. Andreas Neef, Dr. Lorcan Browne, and Dr. Katie Smith for discussions on SGN model development and Antonia Klobe for testing and feed - back on the framework. The speech corpus was provided by Timur Baytukalov (EasyPronounciation.com). The work was funded by the European Research Council (ERC) through the Advanced Grant “OptoHear” under the European Union’s Horizon 2020 Research and Innovation program (grant agreement 670759), the Proof of Concept project “OptoWave” (grant agreement 101113433) and European Innovation Council project “OptoWavePro” (grant agreement 101158920) (to T.M.), the Deutsche Forschungsgemeinschaft (DFG, German Research Foundation) via the Leibniz Program (MO896/5) and the Cluster of Excellence (EXC2067) Multiscale Bioimaging (to T.M.), the Volkswagen Foundation via the SPRUNG program (to T.M.), and the Else Kröner Fresenius Foundation via the Else Kröner Fresenius Center for Optogenetic Therapies (to T.M. and L.J.). This work was also supported by the Ernst Jung Prize for Medicine, the Fondation Pour l’Audition (FPA; RD-2020-10 to T.M.), and the Biotechnology and Biological Sciences Research Council (BBSRC; BB/M019322/1 to D.J.).

## Author contributions

L.K., L.J., and TM designed the study. L.K. implemented the framework. P.N. contributed to the development of the HH model of the SGN. D.J. provided the electrophysiological data for the SGN model fitting. L.J. and T.M. supervised the study. L.K., L.J., and T.M. designed the figures. L.K. and L.J. prepared the figures. L.K. and L.J. prepared the first draft of the manuscript. L.K., L.J., and T.M. prepared the manuscript. All authors discussed the results and commented on the manuscript.

## Competing interests

T.M. is a co-founder of OptoGenTech GmbH.

## Supplementary Information

### Example output of sound coding strategy

A comparison of the output of sound processing for electrical and optical stimulation is provided in Supplementary Figure 1.

**Supplementary Figure 1.**
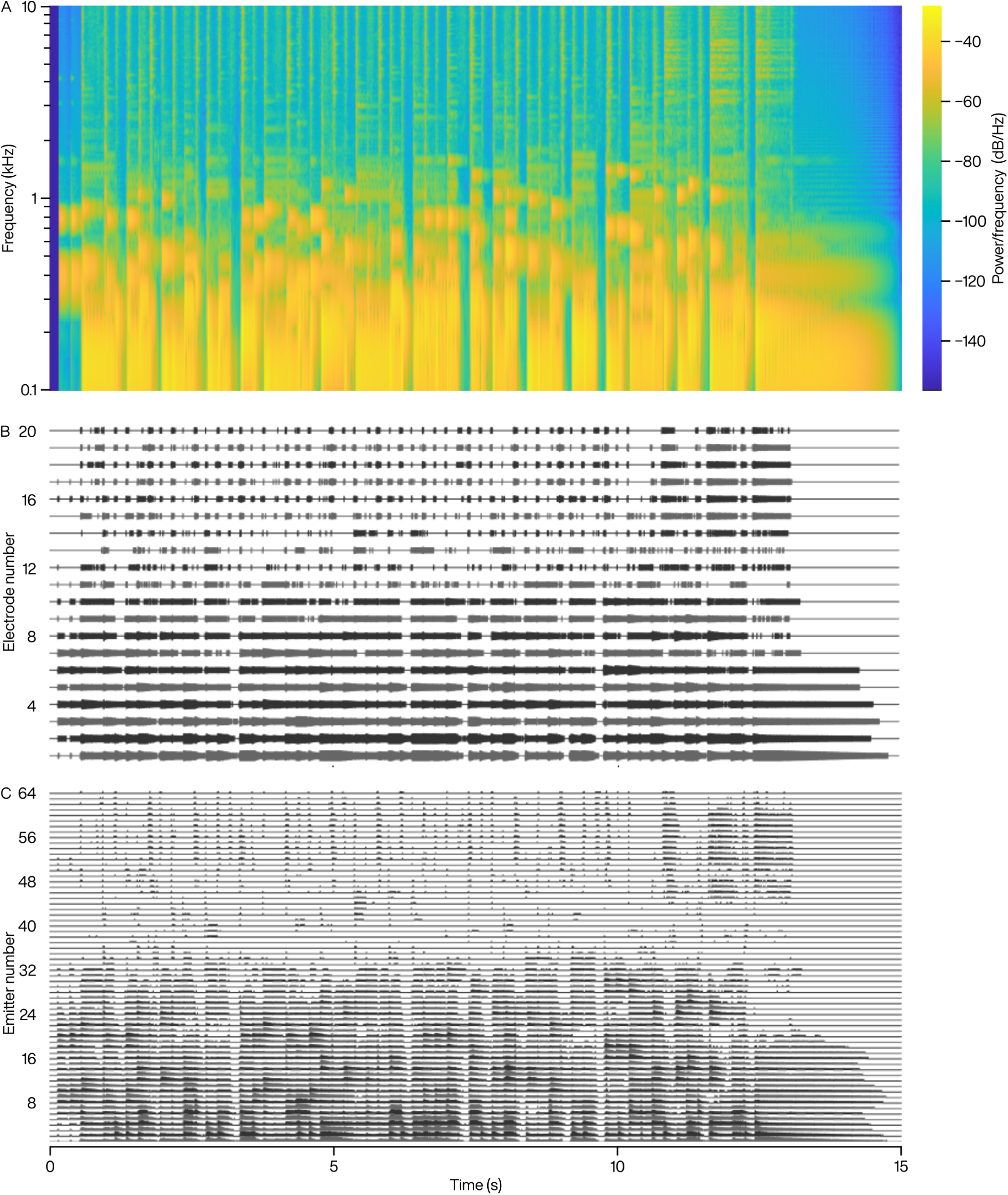
Comparison of electrical and optical sound coding in the framework. (A) A sound spectrogram with a logarithmic scale on the *y*-axis, showcasing the frequency content of a “Happy Birthday” instrumental tune. (B) Electrodogram for the above input with a total of 20 electrodes (*m*) out of which a maximum of 10 were activated in a single frame (*n*) with a pulse rate of 900 pps and 60 μs long biphasic pulses. (C) Emittogram using the same coding strategy but with a set of parameters for optical stimulation, namely, *n*-of-*m* configuration of 32-of-64, 90 pps pulse rate, and 1-ms-long monophasic pulses. Overall, the figure provides a comparison of electrical and optical stimulation in encoding complex auditory stimuli, with a particular emphasis on the potentially enhanced spectral resolution offered by optogenetic cochlear implants.

### Channel models fitting

The channel models are provided in Supplementary Figure 2.

**Supplementary Figure 2.**
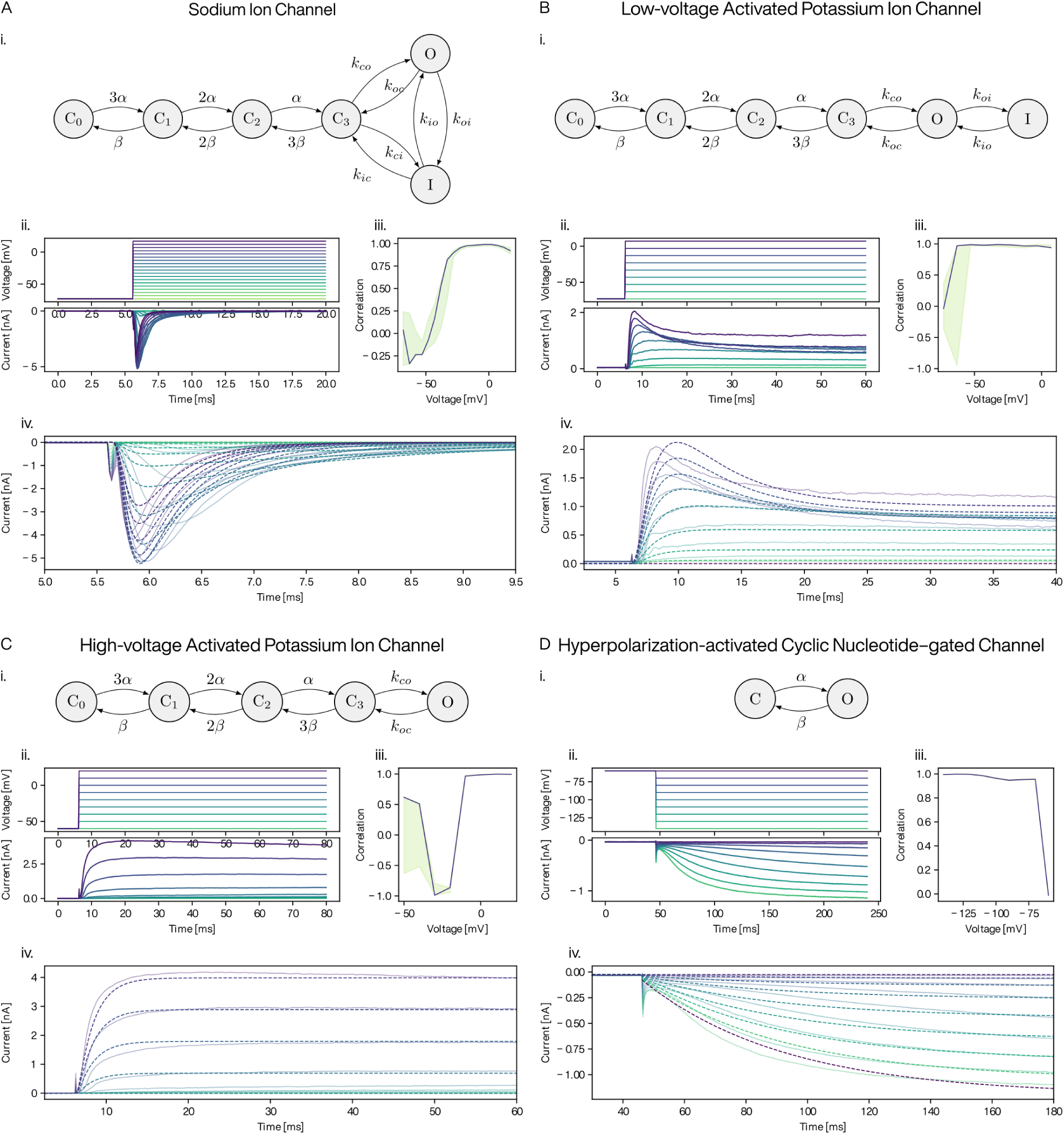
Markov chain models for each ion channel and the fit to the recorded electrophysiological data. (A–D) (i) The ion channel models are based on the Markov chain models described by Tveito and Lines (2016) [32] and Tveito et al. (2016) [62]. (ii) Voltage clamp protocol (top) and average of recorded current traces (bottom). (iii) Correlation between average current traces and the channel model prediction. The light green area shows the minimum and maximum values from all cross-validation batches combined. The dark blue line shows the correlation for the currents estimated with the final model parameters. (iv) Estimated channel currents using final parameters of the model (dashed lines) overlaid on the average of the recorded current traces (full, lighter colored lines) for the same protocols.

### Model parameterization

The rates involved in the model presented in Supplementary Figure 2 are defined by the equations described in Tveito and Lines (2016) [32].

The *Q* _10_ factors, representing the rate of change in channel conductance with a 10°C increase in temperature, and provide a means to calculate channel kinetics at temperatures different from the measured temperature. As the recordings were conducted at room temperature (22–24°C) and we run the model for body temperature (37°C), *Q* _10_ factors were used to multiply the transition rates (Supplementary Table 1).

**Supplementary Table 1.** Q _10_ factors of ion channels implemented in the neuron model.

| Ion channel | $Q_{10}$ factor | Reference |
| --- | --- | --- |
| Voltage-gated sodium channels | 3 | 63 |
| High-voltage activated potassium channels | 3 | 63 |
| Low-voltage activated potassium channels | 3 | 64 |
| Hyperpolarization-activated cyclic nucleotide-gated channels | 4.5 | 65 |
| Chronos | 1.97 (on kinetics)<br>2.92 (off kinetics) | Personal<br>communication with<br>Dr. Andreas Neef<br>(22 February 2023) |

### Computation times

For each block of the framework the computation time was calculated showing that the Hodgkin-Huxley neuron model impacts computation time the most (Supplementary Figure 3).

**Supplementary Figure 3.**
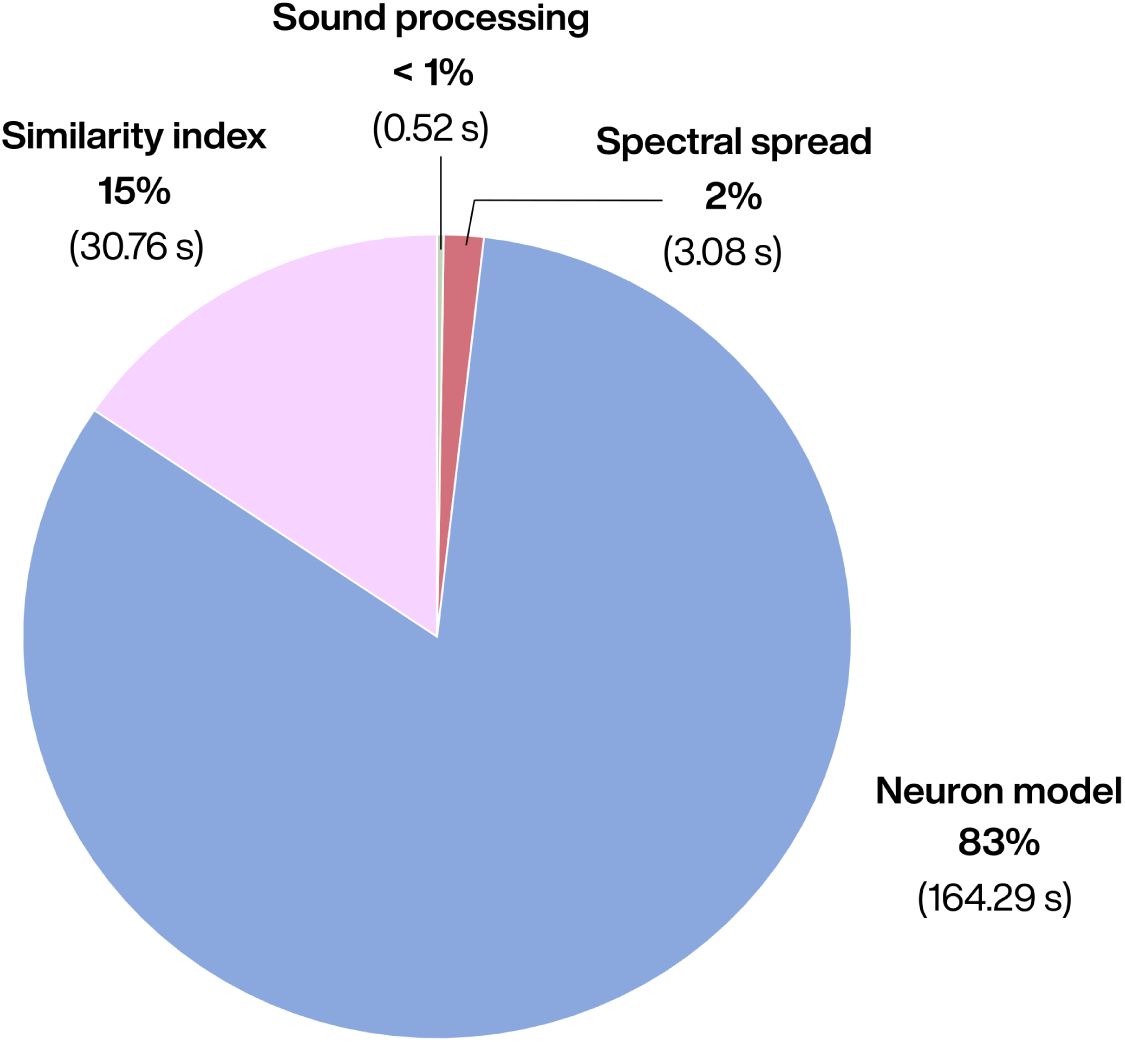
Computation time for each block of the framework. The CPU time was calculated for each block for all the n-of-m configurations of oCI simulations for all audio files. The pie chart shows median values of each block. The Hodgkin-Huxley neuron model utilized the maximum portion of time. The audio files had an average length of 0.79 s and median length of 0.76 s. The computation times were found to be linearly correlated with the length of the input audio (not shown).

## References

1 Micera S, Caleo M, Chisari C, Hummel F C and Pedrocchi A 2020 Advanced Neurotechnologies for the Restoration of Motor Function Neuron 105 604–20

2 Kleinlogel S, Vogl C, Jeschke M, Neef J and Moser T 2020 Emerging Approaches for Restoration of Hearing and Vision Physiol Rev 100 1467–525

3 Zeng F G 2017 Challenges in Improving Cochlear Implant Performance and Accessibility IEEE Trans Biomed Eng 64 1662–4

4 Lenarz T 2017 Cochlear Implant – State of the Art Laryngo-Rhino-Otol 96 S123–51

5 Crowson M G, Semenov Y R, Tucci D L and Niparko J K 2017 Quality of Life and Cost-Effectiveness of Cochlear Implants: A Narrative Review Audiology and Neurotology 22 236–58

6 Fishman K E, Shannon R V and Slattery W H 1997 Speech Recognition as a Function of the Number of Electrodes Used in the SPEAK Cochlear Implant Speech Processor Journal of Speech, Language, and Hearing Research 40 1201–15

7 Friesen L M, Shannon R V, Baskent D and Wang X 2001 Speech recognition in noise as a function of the number of spectral channels: Comparison of acoustic hearing and cochlear implants The Journal of the Acoustical Society of America 110 1150–63

8 Garnham C, O’Driscoll M, Ramsden R and Saeed S 2002 Speech Understanding in Noise with a Med-El COMBI 40+ Cochlear Implant Using Reduced Channel Sets Ear and Hearing 23 540–52

9 Shannon R V, Cruz R J and Galvin J J 2011 Effect of Stimulation Rate on Cochlear Implant Users’ Phoneme, Word and Sentence Recognition in Quiet and in Noise Audiology and Neurotology 16 113–23

10 Berg K A, Noble J H, Dawant B M, Dwyer R T, Labadie R F and Gifford R H 2020 Speech recognition with cochlear implants as a function of the number of channels: Effects of electrode placement The Journal of the Acoustical Society of America 147 3646–56

11 Prevoteau C, Chen S Y and Lalwani A K 2018 Music enjoyment with cochlear implantation Auris Nasus Larynx 45 895–902

12 Limb C J and Roy A T 2014 Technological, biological, and acoustical constraints to music perception in cochlear implant users *Hear*. Res. 308 13–26

13 Hunniford V, Kühler R, Wolf B, Keppeler D, Strenzke N and Moser T 2023 Patient perspectives on the need for improved hearing rehabilitation: A qualitative survey study of German cochlear implant users Frontiers in Neuroscience 17

14 Hernandez V H, Gehrt A, Reuter K, Jing Z, Jeschke M, Mendoza Schulz A, Hoch G, Bartels M, Vogt G, Garnham C W, Yawo H, Fukazawa Y, Augustine G J, Bamberg E, Kügler S, Salditt T, de Hoz L, Strenzke N and Moser T 2014 Optogenetic stimulation of the auditory pathway J. Clin. Invest. 124 1114–29

15 Khurana L, Keppeler D, Jablonski L and Moser T 2022 Model-based prediction of optogenetic sound encoding in the human cochlea by future optical cochlear implants Computational and Structural Biotechnology Journal 20 3621–9

16 Azees A A, Thompson A C, Thomas R, Zhou J, Ruther P, Wise A K, Ajay E A, Garrett D J, Quigley A, Fallon J B and Richardson R T 2023 Spread of activation and interaction between channels with multi-channel optogenetic stimulation in the mouse cochlea Hearing Research 440 108911

17 Dieter A, Duque-Afonso C J, Rankovic V, Jeschke M and Moser T 2019 Near physiological spectral selectivity of cochlear optogenetics Nature Communications 10 1962

18 Dieter A, Klein E, Keppeler D, Jablonski L, Harczos T, Hoch G, Rankovic V, Paul O, Jeschke M, Ruther P and Moser T 2020 μLED-based optical cochlear implants for spectrally selective activation of the auditory nerve EMBO Mol Med e12387

19 Bali B, Lopez de la Morena D, Mittring A, Mager T, Rankovic V, Huet A T and Moser T 2021 Utility of red-light ultrafast optogenetic stimulation of the auditory pathway EMBO Mol Med 13 e13391

20 Mager T, Morena D L de la, Senn V, Schlotte J, D’Errico A, Feldbauer K, Wrobel C, Jung S, Bodensiek K, Rankovic V, Browne L, Huet A, Jüttner J, Wood P G, Letzkus J J, Moser T and Bamberg E 2018 High frequency neural spiking and auditory signaling by ultrafast red-shifted optogenetics Nat Commun 9 1750

21 Keppeler D, Merino R M, Morena D L de la, Bali B, Huet A T, Gehrt A, Wrobel C, Subramanian S, Dombrowski T, Wolf F, Rankovic V, Neef A and Moser T 2018 Ultrafast optogenetic stimulation of the auditory pathway by targeting-optimized Chronos The EMBO Journal 37 e99649

22 Mittring A, Moser T and Huet A T 2023 Graded optogenetic activation of the auditory pathway for hearing restoration Brain Stimul 16 466–83

23 Hanekom T and Hanekom J J 2016 Three-dimensional models of cochlear implants: A review of their development and how they could support management and maintenance of cochlear implant performance Network: Computation in Neural Systems 27 67–106

24 Kalkman R K, Briaire J J and Frijns J H M 2016 Stimulation strategies and electrode design in computational models of the electrically stimulated cochlea: An overview of existing literature Network: Computation in Neural Systems 27 107–34

25 Takanen M, Bruce I C and Seeber B U 2016 Phenomenological modelling of electrically stimulated auditory nerve fibers: A review Network 27 157–85

26 Stadler S and Leijon A 2009 Prediction of Speech Recognition in Cochlear Implant Users by Adapting Auditory Models to Psychophysical Data *EURASIP J. Adv*. Signal Process. 2009 175243

27 Fredelake S and Hohmann V 2012 Factors affecting predicted speech intelligibility with cochlear implants in an auditory model for electrical stimulation Hearing Research 287 76–90

28 Brochier T, Schlittenlacher J, Roberts I, Goehring T, Jiang C, Vickers D and Bance M 2022 From Microphone to Phoneme: An End-to-End Computational Neural Model for Predicting Speech Perception With Cochlear Implants *IEEE Trans*. Biomed. Eng. 69 3300–12

29 Leclère T, Johannesen P T, Wijetillake A, Segovia-Martínez M and Lopez-Poveda E A 2023 A computational modelling framework for assessing information transmission with cochlear implants Hearing Research 432 108744

30 Vandali A E, Whitford L A, Plant K L and Clark G M 2000 Speech perception as a function of electrical stimulation rate: using the Nucleus 24 cochlear implant system Ear Hear. 21 608–24

31 Fromme U 2016 Investigation of voltage- and light-sensitive ion channels (University of Göttingen)

32 Tveito A and Lines G T 2016 A Simple Model of the Sodium Channel Computing Characterizations of Drugs for Ion Channels and Receptors Using Markov Models Lecture Notes in Computational Science and Engineering ed A Tveito and G T Lines (Cham: Springer International Publishing) pp 177–91

33 Smith K E, Browne L, Selwood D L, McAlpine D and Jagger D J 2015 Phosphoinositide Modulation of Heteromeric Kv1 Channels Adjusts Output of Spiral Ganglion Neurons from Hearing Mice J. Neurosci. 35 11221–32

34 Browne L, Smith K E and Jagger D J 2017 Identification of Persistent and Resurgent Sodium Currents in Spiral Ganglion Neurons Cultured from the Mouse Cochlea eNeuro 4 ENEURO.0303-17.2017

35 Gerstner W, Kistler W M, Naud R and Paninski L 2014 Neuronal Dynamics: From Single Neurons to Networks and Models of Cognition (Cambridge: Cambridge University Press)

36 Baytukalov T Audio recordings of words for your app or website

37 Jürgens T, Hohmann V, Büchner A and Nogueira W 2018 The effects of electrical field spatial spread and some cognitive factors on speech-in-noise performance of individual cochlear implant users—A computer model study PLOS ONE 13 e0193842

38 Rutherford M A, Chapochnikov N M and Moser T 2012 Spike Encoding of Neurotransmitter Release Timing by Spiral Ganglion Neurons of the Cochlea J. Neurosci. 32 4773–89

39 Klapoetke N C, Murata Y, Kim S S, Pulver S R, Birdsey-Benson A, Cho Y K, Morimoto T K, Chuong A S, Carpenter E J, Tian Z, Wang J, Xie Y, Yan Z, Zhang Y, Chow B Y, Surek B, Melkonian M, Jayaraman V, Constantine-Paton M, Wong G K-S and Boyden E S 2014 Independent optical excitation of distinct neural populations Nat Methods 11 338–46

40 Hines A and Harte N 2010 Speech intelligibility from image processing Speech Communication 52 736–52

41 Hines A and Harte N 2012 Speech intelligibility prediction using a Neurogram Similarity Index Measure Speech Communication 54 306–20

42 Kandadai S, Hardin J and Creusere C D 2008 Audio quality assessment using the mean structural similarity measure 2008 IEEE International Conference on Acoustics, Speech and Signal Processing 2008 IEEE International Conference on Acoustics, Speech and Signal Processing pp 221–4

43 Gan C, Wang X, Zhu M and Yu X 2011 Audio quality evaluation using frequency structural similarity measure 299–303

44 Fu Q-J and Shannon R V 2000 Effect of stimulation rate on phoneme recognition by Nucleus-22 cochlear implant listeners J. Acoust. Soc. Am. 107 589

45 Plant K, Holden L, Skinner M, Arcaroli J, Whitford L, Law M-A and Nel E 2007 Clinical evaluation of higher stimulation rates in the nucleus research platform 8 system Ear Hear 28 381–93

46 Arora K, Dawson P, Dowell R and Vandali A 2009 Electrical stimulation rate effects on speech perception in cochlear implants International Journal of Audiology 48 561–7

47 Miller C A, Abbas P J, Robinson B K, Nourski K V, Zhang F and Jeng F-C 2006 Electrical excitation of the acoustically sensitive auditory nerve: single-fiber responses to electric pulse trains J Assoc Res Otolaryngol 7 195–210

48 Nicoletti M, Wirtz Chr and Hemmert W 2013 Modeling Sound Localization with Cochlear Implants The Technology of Binaural Listening Modern Acoustics and Signal Processing ed J Blauert (Berlin, Heidelberg: Springer) pp 309–31

49 Mangado N, Pons-Prats J, Coma M, Mistrík P, Piella G, Ceresa M and González Ballester M Á 2018 Computational Evaluation of Cochlear Implant Surgery Outcomes Accounting for Uncertainty and Parameter Variability Frontiers in Physiology 9

50 Kunze K, Goßler C, Reinhardt M, Arnold M, Schwenzer F, Helke C, Reuter D, Keppeler D, Moser T and Schwarz U 2023 Multichannel Laser Diode to Polymer Waveguide Array Coupling with a Double-Aspheric Lens *Appl*. Opt. 62 9353–60

51 Littlefield P D and Richter C-P 2021 Near-infrared stimulation of the auditory nerve: A decade of progress toward an optical cochlear implant Laryngoscope Investigative Otolaryngology 6 310–9

52 Khurana L, Harczos T, Moser T and Jablonski L 2023 En route to sound coding strategies for optical cochlear implants iScience 26 107725

53 Jones M, Warren C, Mashal M, Greenham P and Wyss J 2023 Speech understanding in noise for cochlear implant recipients using a spatial noise reduction setting in an off the ear sound processor with directional microphones Cochlear Implants International 24 311–24

54 Figueiredo J C, Abel S M and Papsin B C 2001 The Effect of the Audallion® BEAMformer Noise Reduction Preprocessor on Sound Localization for Cochlear Implant Users Ear and Hearing 22 539

55 Hersbach A 2014 Noise reduction for cochlear implants

56 Fu Q-J and Nogaki G 2005 Noise Susceptibility of Cochlear Implant Users: The Role of Spectral Resolution and Smearing J Assoc Res Otolaryngol 6 19–27

57 Spoendlin H and Schrott A 1989 Analysis of the human auditory nerve Hearing Research 43 25–38

58 Liberman M C and Oliver M E 1984 Morphometry of intracellularly labeled neurons of the auditory nerve: Correlations with functional properties J. Comp. Neurol. 223 163– 76

59 Moser T, Karagulyan N, Neef J and Jaime Tobón L M 2023 Diversity matters — extending sound intensity coding by inner hair cells via heterogeneous synapses EMBO J 42 e114587

60 Ruggero M A, Santi P A and Rich N C 1982 Type II cochlear ganglion cells in the chinchilla Hearing Research 8 339–56

61 Spoendlin H 1972 Innervation densities of the cochlea Acta Otolaryngol. 73 235–48

62 Tveito A, Lines G T, Edwards A G and McCulloch A 2016 Computing rates of Markov models of voltage-gated ion channels by inverting partial differential equations governing the probability density functions of the conducting and non-conducting states Math Biosci 277 126–35

63 Schwarz J R and Eikhof G 1987 Na currents and action potentials in rat myelinated nerve fibres at 20 and 37° C Pflugers Arch. 409 569–77

64 Hille B 2001 Ion channels of excitable membranes (Sunderland Massachusetts USA: Sinauer Associates)

65 Negm M 2008 An Improved Stochastic Hodgkin-Huxley Based Model of a Node of Ranvier for Cochlear Implant Stimulation Thesis

